# Homologous recombination-based genome editing by clade F AAVs is inefficient in the absence of a targeted DNA break

**DOI:** 10.1101/704197

**Authors:** Geoffrey L. Rogers, Hsu-Yu Chen, Heidy Morales, Paula M. Cannon

## Abstract

Adeno-associated virus (AAV) vectors are frequently used as donor templates for genome editing by homologous recombination. Although modification rates are typically under 1%, they are greatly enhanced by targeted double-stranded DNA breaks (DSBs). A recent report described clade F AAVs mediating high-efficiency homologous recombination-based editing in the absence of DSBs. The clade F vectors included AAV9 and a series isolated from human hematopoietic stem/progenitor cells (HSPCs). We evaluated these vectors by packaging homology donors into AAV9 and an AAVHSC capsid and examining their ability to insert GFP at the CCR5 or AAVS1 loci in human HSPCs and cell lines. As a control we used AAV6, which effectively edits HSPCs, but only when combined with a targeted DSB. Each AAV vector promoted GFP insertion in the presence of matched CCR5 or AAVS1 zinc finger nucleases (ZFNs), but none supported detectable editing in the absence of the nucleases. Rates of editing with ZFNs correlated with transduction efficiencies for each vector, implying no differences in the ability of donor sequences delivered by the different vectors to direct genome editing. Our results therefore do not support that clade F AAVs can perform high efficiency genome editing in the absence of a DSB.

## Introduction

Interest in the use of genome editing technologies to correct human genetic mutations or precisely insert therapeutic gene cassettes at defined loci has greatly increased in recent years.^1^ In the most common approach, targeted nucleases such as CRISPR/Cas9, zinc finger nucleases (ZFNs) or transcription activator-like effector nucleases (TALENs) generate double-stranded DNA breaks (DSB) at a specific sequence in the genome with high precision.^2–4^ The subsequent repair of DSBs can result in insertions and deletions (indels) from the activity of the non-homologous end joining (NHEJ) DNA repair pathway, and this can be leveraged to disrupt an open reading frame or genetic element.^5, 6^ In contrast, by also introducing a DNA donor template that is homologous to the chromosomal DNA surrounding the DSB, it is possible to harness homology-directed repair (HDR) pathways and thereby engineer specific DNA changes into the host genome.^7^

DNA homology donors can be provided by plasmids, oligonucleotides or viral genomes.^4, 5, 8-10^ Among these, adeno-associated viral (AAV) vectors have emerged as particularly effective vehicles for delivery of DNA homology donors.^5, 11-13^ AAV is a small parvovirus encapsidating a single-stranded DNA genome of about 4.7 kb.^14^ Recombinant AAV vector genomes contain only the viral inverted terminal repeats (ITRs) and persist as stable episomal DNA, with expression detected for over a decade *in vivo.*^15–18^ In addition, many different AAV serotypes are available, comprising both natural and engineered capsids, that allow transduction of a wide variety of cell types and tissues *in vitro* and *in vivo*.^19–22^ Accordingly, AAV vectors are being evaluated as gene delivery vectors in a number of human clinical trials.^22, 23^

AAV vectors have a long history of use as homology donors in genome targeting applications. Such studies over the past 20 years have typically reported gene insertion rates below 1% in the absence of a targeted nuclease.^24–27^ In contrast, combining AAV donors with targeted nucleases has led to high rates of genome editing, most notably in hematopoietic cells, where combining AAV serotype 6 donors with ZFNs,^5, 11^ TALENs,^13^ or CRISPR/Cas9^12, 28^ has resulted in gene editing rates of 15-60% in T cells, B cells and CD34^+^ hematopoietic stem and progenitor cells (HSPCs). Nuclease-mediated engineering is accompanied by potential risks from off-target DNA breaks, although improvements in nuclease engineering^29–32^ and enhanced off-target detection methods^33–35^ are reducing these concerns. At the same time, editing technologies that do not require DSBs are also being developed, such as those that exploit base editing^36–38^ and potentially transposon integration.^39^

Recently, *Smith et al.*^40^ reported that the use of AAV vectors generated with capsids from clade F viruses could mediate highly efficient HDR in the absence of targeted DNA breaks. Clade F includes serotype AAV9, as well as a closely related family of novel capsids termed AAVHSCs that were previously isolated by their group from human HSPCs.^41^ These clade F AAV capsids were reported to mediate stable gene insertion in both cell lines and primary human HSPCs, resulting in scarless modifications that are consistent with HDR in upwards of 50% of HSPCs at the *AAVS1* locus and 8% of HSPCs at two sites in the *IL2RG* gene. Such an effect was not seen with identical AAV genomes packaged into capsids from other serotypes. The authors further reported that this genome editing required BRCA2, suggesting dependence on homologous recombination, although a more thorough explanation of the underlying mechanism was not obtained. Given the departure of these findings from previous studies of nuclease-free genome editing using AAV vectors, we were interested to evaluate the utility of clade F AAV vectors in genome editing, both with and without matched targeted nucleases.

## Results

### Transduction of human HSPCs and cell lines by AAV6 and clade F vectors

Smith *et al.* reported that AAV genomes packaged into clade F capsids could direct high levels of homologous recombination-based genome editing in the absence of a catalyzing DSB.^40^ The authors further suggested that the concentration of AAV genomic DNA within the nucleus may have played a role in this process, since they reported significantly better transduction and nuclear entry rates in human CD34^+^ HSPCs when using clade F AAVs compared to other serotypes, including AAV6, and that the efficiency of transduction correlated with the editing frequencies obtained. These results were somewhat surprising, given that we and others have previously reported that AAV6 vectors more efficiently transduce HSPCs than the prototype clade F vector, AAV9.^5, 42^ We therefore directly compared the ability of AAV6 and AAV9 vectors to transduce both HSPCs and three different cell lines. Using an AAV genome carrying only a CMV-GFP cassette and no homology arms, we observed transduction rates of HSPCs after 2 days that ranged from ~20-80% when using AAV6 at multiplicities of infection (MOIs) between 10^3^ and 10^5^. In contrast, the matched AAV9 vectors transduced less than 3% of cells at the highest MOI of 10^5^ (Fig. 1A). Similarly, we found that AAV6 vectors transduced all three cell lines better than the AAV9 vectors (Fig. 1B). This panel included the K562 cell line that Smith *et al*. reported to be efficiently edited by AAV9 vectors in the absence of a targeted nuclease. In contrast, we observed the highest rates of transduction using AAV9 vectors on HEK-293T cells, which were reported to be poorly edited by several clade F AAVHSCs by Smith *et al.* Regardless of the initial levels of transduction, GFP expression in the HEK-293T populations declined greatly for both AAV6 and AAV9 vectors between days 2 and 17, consistent with the dilution of unintegrated AAV genomes in a population of dividing cells (Fig. 1C). This observation was consistent with expectations for a vector genome containing no homology sequences and suggested that the persistence of GFP expression over time could be used to detect stable gene insertion.

**Figure 1.**
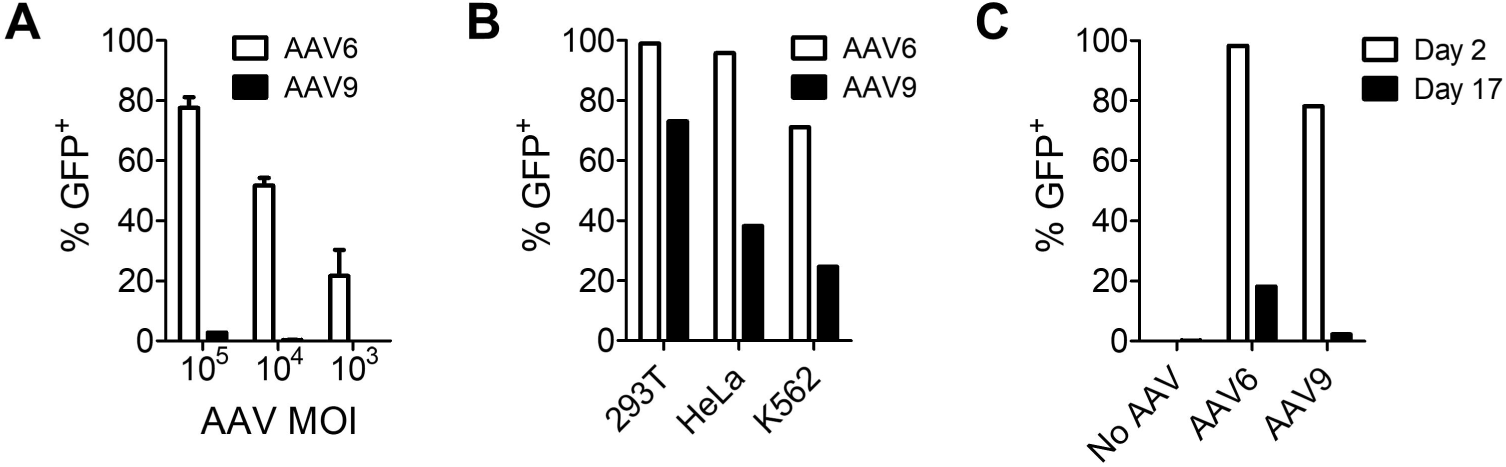
Transduction of human HSPCs and cell lines by AAV6 and AAV9. (A) HSPCs were transduced with AAV6 or AAV9 vectors containing a CMV-GFP cassette at indicated MOIs and GFP expression was measured 2 days later by flow cytometry, for *n* = 2 HSPC donors. Data are shown as mean ± SEM. (B) HEK-293T, HeLa and K562 cells were transduced with AAV6 or AAV9 CMV-GFP vectors at an MOI of 10^5^, and GFP expression was measured 2 days later by flow cytometry. (C) HEK-293T cells were transduced with AAV6 or AAV9 CMV-GFP vectors at an MOI of 5 × 10^5^, and GFP expression was measured at day 2 and day 17 by flow cytometry.

### AAV vectors mediate efficient homology-directed gene insertion in K562 cells only in the presence of a targeted nuclease

To further investigate targeted integration by clade F AAVs we generated a mutant capsid, AAV9-G505R. This vector matches the reported amino acid sequence of AAVHSC13 and AAVHSC17 and contains a point mutation (G505R) that is conserved among 6 of the 17 AAVHSC serotypes described.^41^ We produced AAV6, AAV9 and AAV9-G505R vectors packaging a genome comprising a PGK-GFP cassette flanked by CCR5 homology arms (Fig. 2A), with which we have previously evaluated nuclease-driven HDR-mediated gene editing in HSPCs.^5^ K562 cells transduced with AAV6, AAV9, or AAV9-G505R vectors were 48.0%, 8.81%, and 4.38% GFP^+^ respectively after 2 days, indicating that all three vectors were able to transduce these cells at varying efficiencies (Fig. 2B). After 14 days, all cell populations were <0.5% GFP^+^ suggesting that the vast majority of the initial GFP expression at day 2 originated from episomal AAV genomes and that minimal, if any, integration of the transgene cassette had occurred (Fig. 2B-C). In contrast, when the cells were also electroporated with mRNA coding for CCR5-specific ZFNs, stable GFP expression was detected after 14 days at frequencies that were similar to the day 2 transduction values (Fig 2B-C). Site-specific insertion of the GFP cassette at the CCR5 locus was confirmed using an in-out PCR assay^5^ which only produced a signal in the ZFN-treated samples (Fig. 2D). Together, these observations indicate that although each of the three AAV vectors could deliver homology donors that promote HDR-mediated gene addition in the presence of a targeted DNA break, they were unable to do so efficiently in the absence of such a DSB.

**Figure 2.**
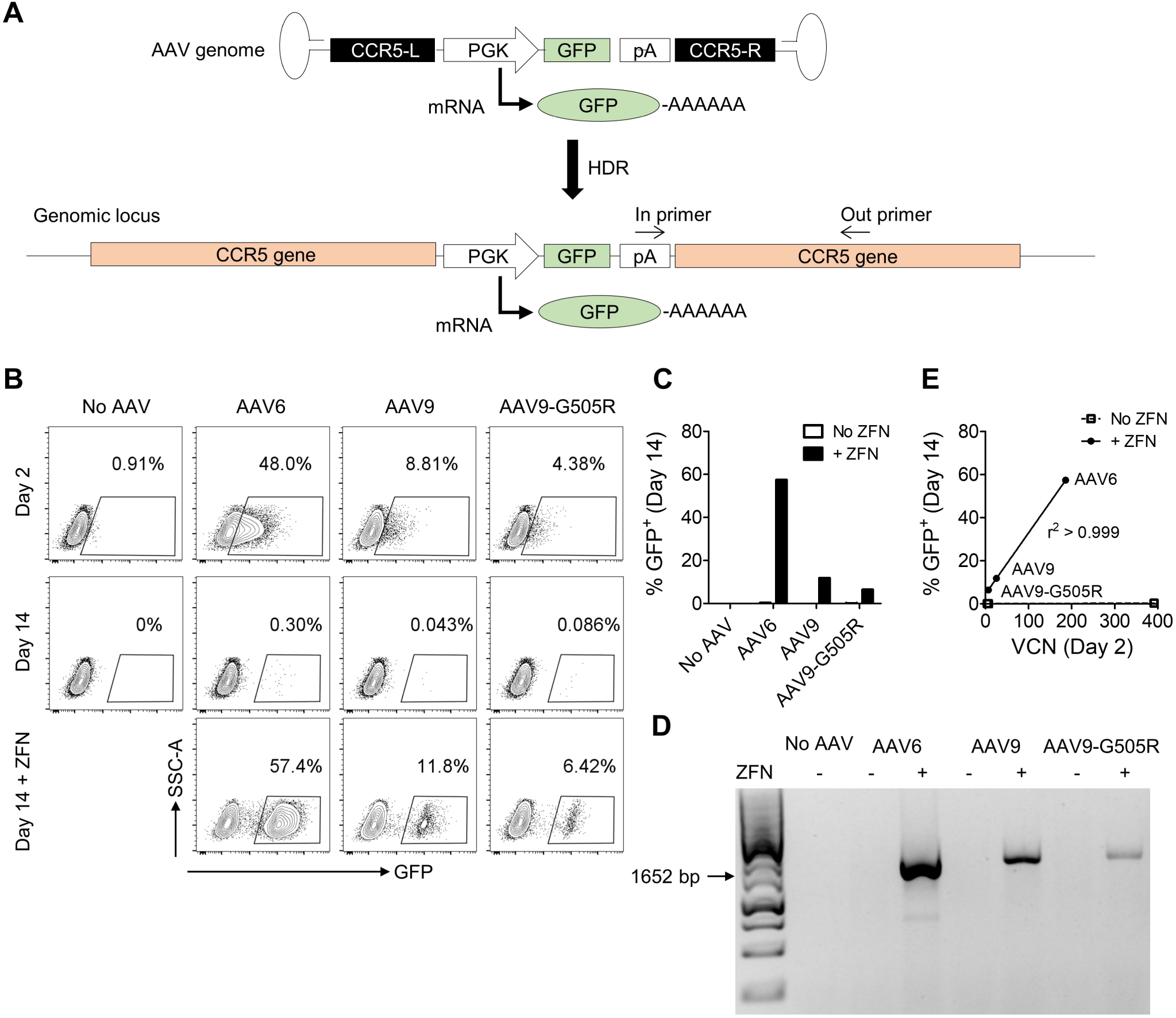
Genome editing K562 cells at the CCR5 locus requires a targeted nuclease. (A) Schematic of the CCR5-PGK-GFP AAV vector genome and the result of HDR genome editing at the CCR5 locus. PGK promoter-driven GFP expression can occur from episomal AAV genomes or following insertion into the host DNA. The position of in-out PCR primers, to detect site-specific insertion, are indicated. (B) K562 cells were transduced with AAV6, AAV9, or AAV9-G505R vectors packaging CCR5-PGK-GFP genomes at an MOI of 10^4^, and samples were electroporated with CCR5-specific ZFN mRNA as indicated. Flow cytometry plots are shown for indicated treatments and times. (C) Comparison of GFP expression after 14 days in indicated samples, treated with or without ZFN mRNA. (D) In-out PCR to detect insertion of the PGK-GFP cassette at CCR5. (E) Correlation between cell-associated vector copy number (VCN) at day 2 versus GFP expression by flow cytometry at day 14, for indicated samples.

Since the transduction rates at day 2 seemed to be predictive of the genome editing rates achieved in the presence of the ZFNs, we performed AAV vector genome copy number analysis at day 2 by ddPCR. The number of cell-associated AAV genomes at day 2 strongly correlated with the frequency of GFP^+^ cells at day 14 (r^2^ > 0.999) for all 3 serotypes when co-delivered with the ZFN mRNA (Fig. 2E). These data suggest that transduction efficiency rather than any intrinsic properties of the different AAV serotypes governs the rate of nuclease-dependent gene addition.

To verify that the lack of nuclease-independent genome editing we observed for AAV6 and the two clade F vectors was not specific to the CCR5 locus, we generated AAV6, AAV9, and AAV9-G505R vectors with genomes that targeted the AAVS1 locus. To mimic the strategy used by Smith *et al.*, the vector genomes comprised a promoter-less GFP cassette whose expression relied on an upstream splice acceptor and T2A sequence, and containing AAVS1 homology arms (AAVS1-SA-2A-GFP) (Fig. 3A).^40^ Since this construct lacks an internal promoter, GFP expression should only be observed following insertion into the host genome downstream of a promoter and within an intron and, as expected, episomal expression was not detected at day 2 in any of the samples transduced with AAV only (Fig 3B). At day 14, GFP expression (Fig. 3B-C) and site-specific insertion (Fig. 3D) in K562 cells was only observable above background levels in cells also treated with AAVS1-specific ZFN mRNA, with frequencies that were comparable to those observed at CCR5. Vector copy number at day 2 also strongly correlated with GFP expression at day 14 in the presence of ZFNs, also confirming the importance of transduction efficiency in determining the rates of nuclease-dependent gene editing at this locus (Fig. 3E).

**Figure 3.**
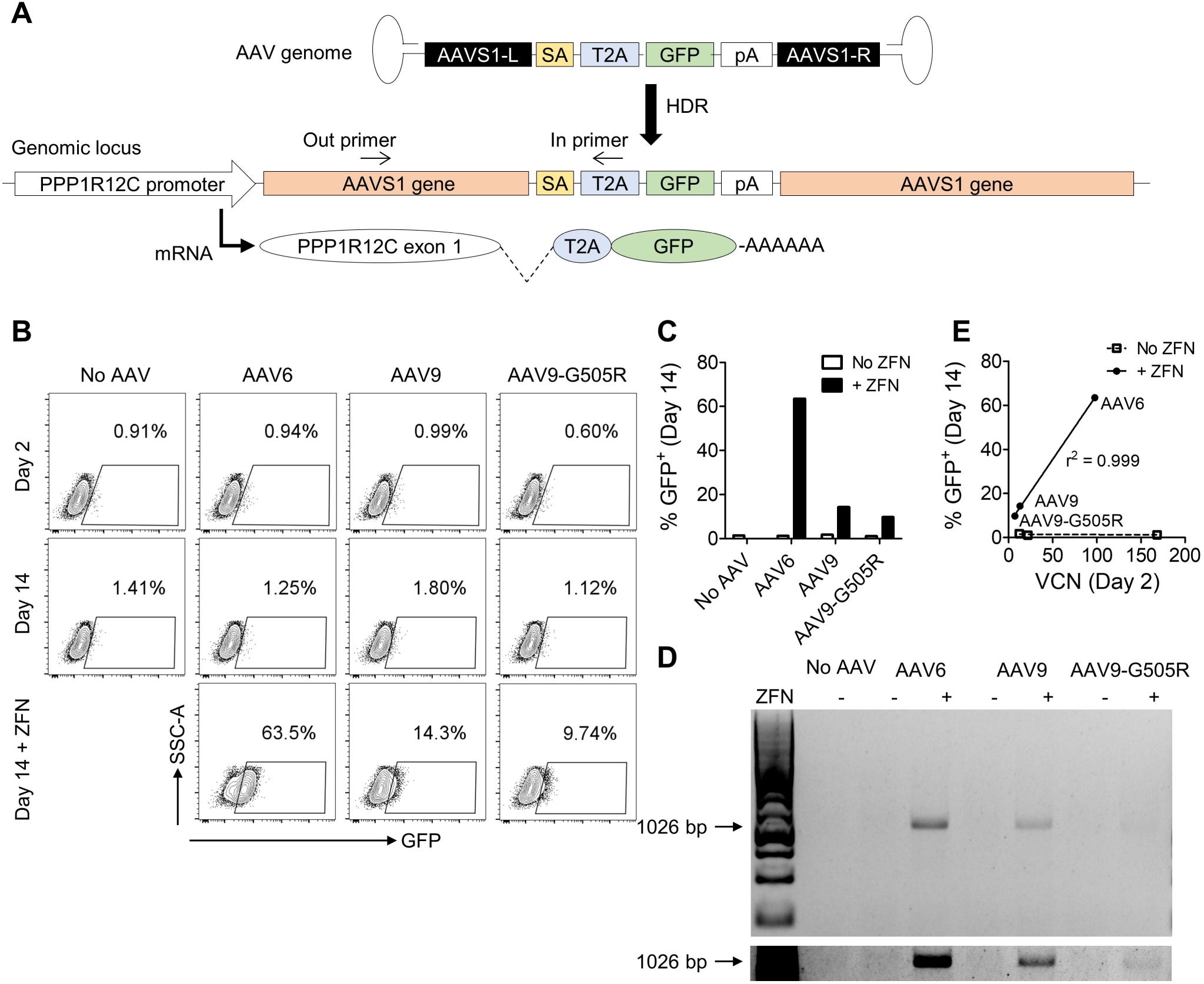
Genome editing K562 cells at the AAVS1 locus requires a targeted nuclease. (A) Schematic of the AAVS1-SA-2A-GFP AAV vector genome and the result of HDR genome editing at the AAVS1 locus. GFP expression occurs following site-specific integration and splicing of the edited *PPP1R12C* transcript. The position of in-out PCR primers, to detect site-specific insertion, are indicated. (B) K562 cells were transduced with AAV6, AAV9, or AAV9-G505R vectors packaging AAVS1-SA-2A-GFP genomes at an MOI of 10^4^, and samples were electroporated with AAVS1-specific ZFN mRNA as indicated. Flow cytometry plots are shown at indicated treatments and times. (C) Comparison of GFP expression after 14 days in indicated samples, treated with or without ZFN mRNA. (D) In-out PCR to detect insertion of the PGK-GFP cassette at AAVS1. The lower panel is a cropped overexposure of the same gel with increased brightness to improve visualization of the AAV9-G505R samples. (E) Correlation between cell-associated vector copy number (VCN) at day 2 versus GFP expression by flow cytometry at day 14, for indicated samples.

### Efficient transduction does not support nuclease-independent genome editing by clade F AAVs in HEK-293T cells

A potential limitation of our analyses in K562 cells is the relatively inefficient transduction (<10%) we observed with clade F AAVs. To address this concern, we next tested the panel of vectors in HEK-293T cells, which were more permissive to AAV9 transduction in our hands (Fig. 1B). Using a higher MOI of 5 × 10^5^ of the CCR5-PGK-GFP vectors, we observed GFP expression at day 2 in 97.5% of cells with AAV6, 55.9% with AAV9, and 19.8% with AAV9-G505R (Fig. 4A). Despite these high initial rates of AAV transduction, GFP expression declined significantly in all populations by day 17, mirroring the drop we previously observed using AAV vectors lacking homology arms (Fig. 1C), and again suggesting that the GFP expression cassette had not been incorporated into the genome. Similarly, cells transduced with equivalent MOIs of AAV vectors packaging the AAVS1-SA-2A-GFP genome did not exhibit GFP expression above background at either day 2 or day 17, providing further evidence that site-specific gene editing had not occurred (Fig. 4B). Together, these results suggest that neither AAV6, AAV9, nor AAV9-G505R vectors can mediate efficient genome editing in the absence of a targeted nuclease, even when high levels of vector transduction have occurred.

**Figure 4.**
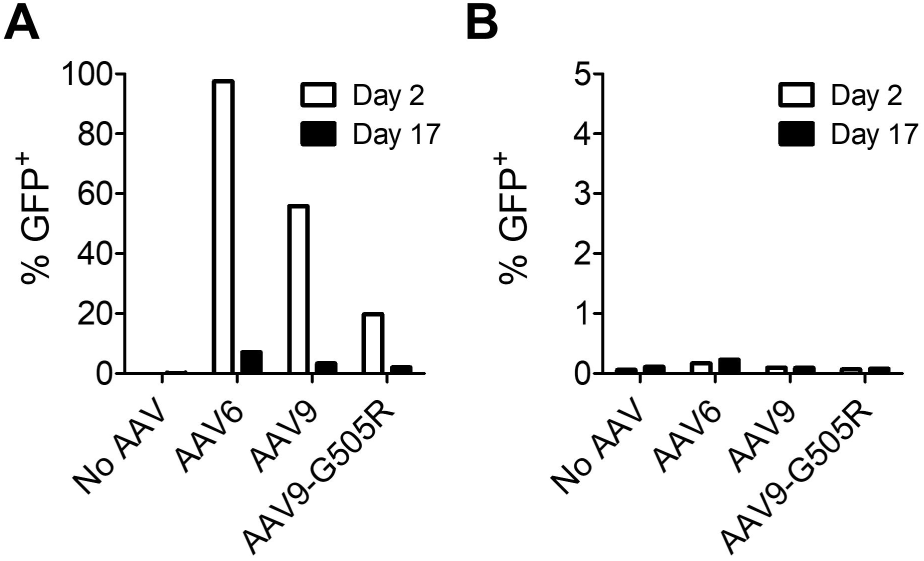
Lack of nuclease-free genome editing in HEK-293T cells. HEK-293T cells were transduced with AAV6, AAV9, or AAV9-G505R vectors packaging either (A) CCR5-PGK-GFP genomes or (B) AAVS1-SA-2A-GFP genomes, at an MOI of 5 × 10^5^, and GFP expression was measured at indicated timepoints by flow cytometry.

### Lack of homology-directed gene insertion in HSPCs by AAV vectors in the absence of a matched targeted nuclease

Finally, we investigated the editing efficiency of AAV homology donors in primary human HSPCs. In agreement with our data obtained using the CMV-GFP reporter vectors (Fig. 1A), GFP levels at day 2 post-transduction for CCR5-PGK-GFP constructs were much more efficient when packaged in AAV6 capsids than either AAV9 or AAV9-G505R (Fig. 5A). Indeed, the clade F vectors only produced 0.54% and 0.32% GFP^+^ cells, respectively, for AAV9 and AAV9-G505R when using an MOI of 5 × 10^4^. This MOI is comparable to that employed by Smith *et al.* in HSPCs.^40^ By day 10, GFP expression from all of the cells receiving AAV vectors alone had fallen to below 0.3% GFP^+^ (Fig. 5B-C). In contrast, combining the vectors with CCR5-specific ZFNs promoted stable GFP expression at day 10 that was above the frequencies observed for the vector only controls (Fig. 5B-C). The levels obtained with the two clade F vectors were <1%, but this likely reflected their poor ability to transduce HSPCs, which would limit genome editing rates. Finally, we also evaluated stable GFP expression levels when AAVS1-specific ZFNs were combined with the CCR5 homology donor vectors (Fig. 5B-C). This mismatched nuclease will introduce a DSB at a non-homologous locus, which can lead to low levels of NHEJ-mediated end capture of AAV genomes,^5^ and indeed we observed a slight enhancement of GFP expression for the AAV6 vectors above the background levels with no ZFN.

**Figure 5.**
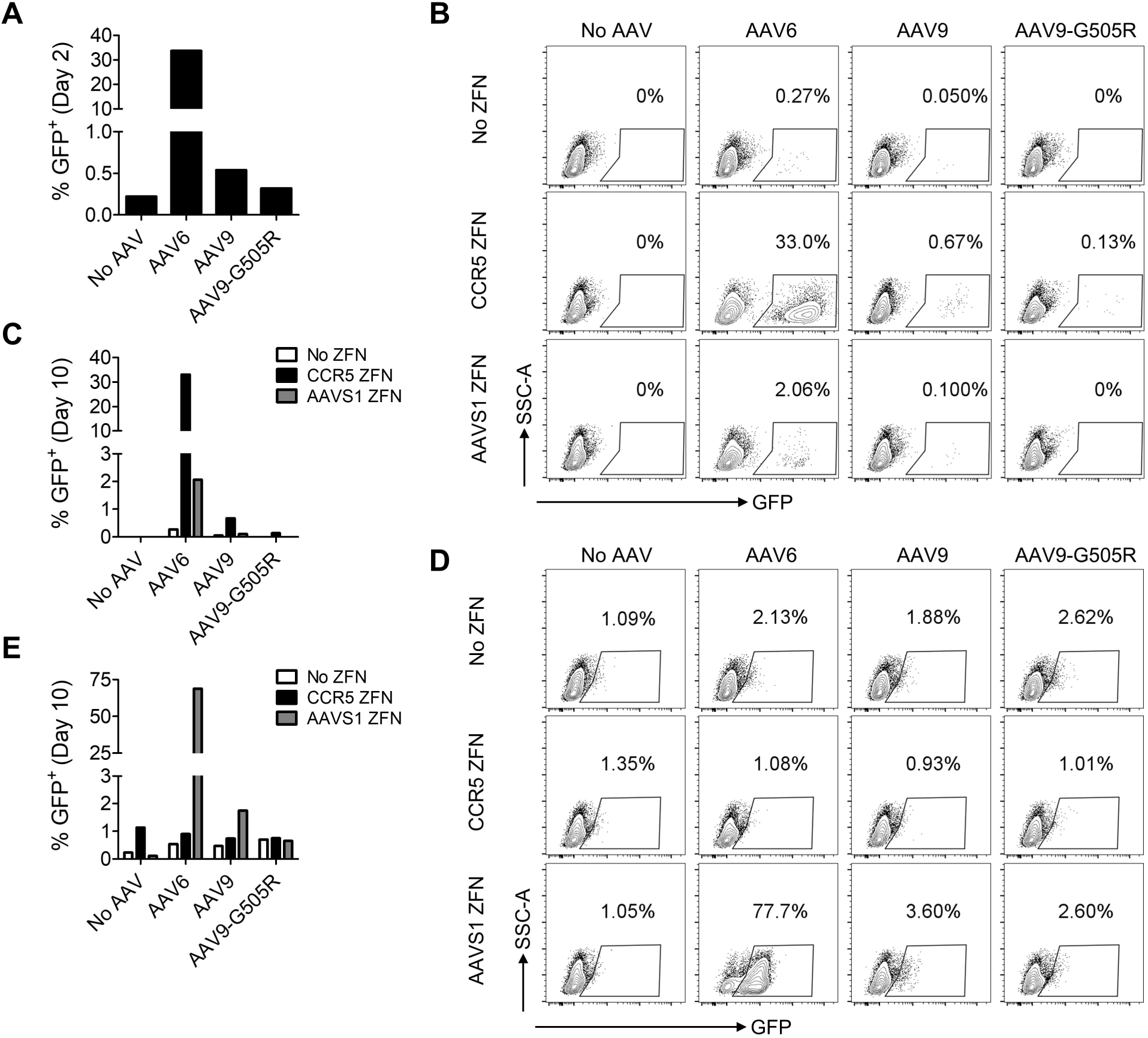
Genome editing in HSPCs by AAV6 and clade F vectors requires matched nucleases. HSPCs were transduced with AAV6, AAV9, or AAV9-G505R vectors packaging (A-C) CCR5-PGK-GFP vector genomes, or (D-E) AAVS1-SA-2A-GFP vector genomes, at an MOI of 5 × 10^4^. Samples were electroporated with CCR5- or AAVS1-specific ZFN mRNAs as indicated. GFP expression was measured at day 2 or day 10 post-electroporation by flow cytometry. (A) Graphs of day 2 GFP measurements from AAV vector only samples show rates of initial AAV transduction. (B,D) Flow cytometry analyses for indicated treatments at day 10. (C, E) Graphical representation of stable GFP expression at day 10.

We repeated these analyses using vectors containing the AAVS1-SA-2A-GFP homology donor genome, delivered alone to HSPCs or in combination with matched (AAVS1) or mismatched (CCR5) ZFNs (Fig. 5D-E). The results mirrored those seen at the CCR5 locus, since GFP expression above background was only detected in cells also receiving the AAVS1-specific ZFNs. A comparable hierarchy of efficiency was observed among the 3 AAV capsids, though even with the AAVS1 ZFNs, GFP expression was not detectable above background for AAV9-G505R vectors, presumably due to its low initial transduction rate. In all cases, no GFP expression was observed in the presence of the mismatched CCR5 ZFNs, despite the possibility of end-capture events, but this would be expected given the lack of an internal promoter in the vector genome.

## Discussion

The hijacking of homology-directed repair pathways to introduce precise changes into a cell’s genome is a central aspect of many genome editing technologies. Although HDR-mediated editing only requires a homologous DNA template, the process is very inefficient on its own.^43^ Consequently, the discovery that targeted DSBs could catalyze HDR editing was highly significant,^44^ with the subsequent development of targeted nucleases providing the necessary technological advancement to reach the current capabilities of genome editing. Despite these successes, targeted nucleases and the creation of DSBs raise concerns about genotoxicity and off-target effects, so that approaches that enhanced the efficiency of nuclease-free HDR editing would be welcome.

DNA homology templates can be provided by plasmids, single-stranded oligonucleotides or the genomes of DNA viruses. In the absence of a DSB, AAV is often used to promote HDR editing and frequencies up to 1% have been reported.^24–27^ The advantages of AAV in this process are unclear, although its stable episomal genome, the availability of single-stranded and double-stranded forms of the genome, and the structure of the terminal ITR sequences have been suggested as factors. In addition, the wide variety of natural and engineered capsid serotypes of AAV allow the delivery of DNA templates to a multitude of cell types, both *in vitro* and *in vivo*. Identification of AAV6 as having good tropism for human HSPCs was central to our own efforts to develop nuclease-mediated genome editing in these cells.^5^

AAV vectors are also used in more conventional gene therapy approaches and are accruing a record of safety in human clinical trials. Random integration into the host genome reportedly occurs at extremely low frequencies (<0.1%) and appears to exhibit no deleterious preferences in targeting.^45^ Some studies have suggested a link between AAV delivery and hepatocellular carcinoma in mice, though others did not reproduce these findings in nonhuman primates and patients.^46–48^ To date, no reports have indicated germline modification. However, in the absence of a targeted DSB, the ability of AAV vectors to specifically modify the human genome has been much more limited, making therapeutic applications of AAV mediated HDR more challenging and potentially dependent on *in vivo* selection or expansion.^25, 26, 49^ To achieve targeted integration *in vivo*, clinical trials are now progressing using AAV vectors to deliver both genome editing components and the donor DNA for MPS I (NCT02702115), MPS II, (NCT03041324) and hemophilia B (NCT02695160), though the mechanism of gene insertion can include both HDR and site-specific NHEJ-mediated insertion of the AAV transgene.^50, 51^ However, the requirement for both a nuclease and a donor template complicates this *in vivo* delivery platform. Hence, the report of efficient homologous recombination by clade F AAV vectors in the absence of targeted nucleases is a potentially exciting discovery.^40^

To evaluate this reported ability of clade F vectors, we packaged homology templates for the CCR5 or AAVS1 loci into AAV9 vectors, or those containing an additional point mutation G505R, which corresponds to the reported variants AAVHSC13 and AAVHSC17.^41^ As a control, we also included AAV6 vectors. The AAVS1 homology donor genome we designed mimicked the approach used by Smith *et al.*,^40^ where a promoter-less GFP reporter was placed downstream of a splice acceptor cassette, so that site-specific integration into the *PPP1R12C* intron could result in GFP expression. In contrast, the CCR5 homology construct contained an internal PGK promoter, expected to drive GFP expression from both episomal and integrated forms of the vector genome. We confirmed that both donor templates were HDR-competent by demonstrating stable GFP expression in the presence of a matched ZFN and further confirmed that integration had occurred into the targeted loci using a specific in-out PCR. However, in the absence of the matched nucleases, we observed only background levels of GFP expression and no detectable PCR products when all 6 of the vectors were tested in human HSPC and two different cell lines.

A possible explanation proposed for the properties of the clade F vectors described by Smith *et al*. was enhanced delivery to the nuclei of target cells. Although we did not detect nuclease-independent editing, we consistently observed a strong correlation between the AAV transduction rates we measured on day 2 and the genome editing frequency we measured at later time points in the presence of matched ZFNs. This observation was particularly compelling in K562 cells, where the number of cell-associated AAV genomes nearly perfectly correlated with the rate of stable GFP insertion that was achieved across the three capsids. Similarly, the GFP transduction pattern we observed in HSPCs at day 2 matched the relative nuclease-mediated editing rates that persisted at day 10 (AAV6 >>> AAV9 > AAV9-G505R). Together, these data suggest that our clade F vectors did not display a unique propensity for HDR; instead, the concentration of homology donor template delivered to the cell was the major determinant of editing rates for all three AAV serotypes tested.

Other aspects of our work did not agree with the results reported by Smith *et al*., and it is possible that these could account for the large discrepancies we observed in nuclease-free editing capabilities. For example, the relative tropism we obtained with AAV6 and AAV9 vectors on HSPCs is in stark contrast to their observations. In HSPCs and in cell lines, we observed superior transduction using AAV6 over either of the clade F vectors we tested at equivalent MOIs, which aligns with previously published comparisons of AAV6 and AAV9 tropism in HSPCs from both our group and others.^5, 42^ Indeed, AAV6 is now routinely used by many different labs to transduce HSPCs for genome editing applications.^12, 13, 52^ Our observations are also consistent with reports that have described overall low transduction *in vitro* with AAV9.^20, 21^ In contrast, Smith *et al.* report 27-fold higher nuclear entry in HSPCs for a pool of clade F vectors compared to AAV6, whereas we observed roughly the inverse in K562 cells–*i.e.* an average of 36- and 48-fold higher VCNs with AAV6 compared to AAV9 and AAV9-G505R, respectively.^40^

It is also possible that the published capsid sequence for AAVHSC13 and AAVHSC17^41^ is not sufficient to reproduce these results. These variants differ from AAV9 by a single G505R mutation in VP3, although AAVHSC17 was also reported to contain a silent mutation in VP1.^41^ Confusingly, both transduction experiments and genome editing results in the recent Smith *et al.* report give differing efficiencies for AAVHSC13 and AAVHSC17.^40^ It is unclear why these two capsids would behave differently, unless the silent mutation in AAVHSC17 has some unusual regulatory function, such as altering the traditional 1:1:10 VP1:VP2:VP3 ratio of AAV capsid proteins. Nevertheless, these potential subtleties of AAVHSC capsids do not explain the different results obtained with the prototype clade F member AAV9, which is a well-studied serotype that is used in many applications.

Although we cannot explain these observed *in vitro* tropism differences, we recognized that the lower transduction efficiencies we obtained when using clade F vectors compared to AAV6 were a confounding factor. We achieved less than 10% GFP^+^ cells in K562 cells and less than 1% GFP^+^ cells in HSPCs, even when using an MOI that was 16-fold higher than we typically need to achieve HDR insertion rates in HSPCs at the CCR5 locus of 15-50% (5 × 10^4^ vs. 3 × 10^3^) (unpublished observations).^5^ While it is possible that our transduction levels were too low to detect nuclease-free HDR editing in the assays we used, the MOIs we used were within the range used by Smith *et al*,^40^ although the well-documented potential variability of AAV titers between labs must be considered when making these sorts of comparisons.^53, 54^ Additionally, the vector copy numbers we found in K562 cells were at least as high as those reported by Smith in HSPCs, albeit with the opposite observations of relative efficiency between AAV6 and the clade F AAVs. These data suggest that we are using a similar range of AAV doses.

Nevertheless, to try to rule out such a dose effect, we repeated the analyses in HEK-293T cells, where we could achieve transduction rates of >50% of cells with AAV9 vectors, and >20% with AAV9-G505R. These transduction frequencies are in line with those reported by Smith *et al*. to yield gene insertion rates in excess of 10%, depending on the specific clade F serotype used. However even with this high level of transduction in HEK-293T cells, we found no evidence of efficient site-specific genome editing in the absence of a targeted nuclease with AAV9 or AAV9-G505R vectors. Possibly, higher MOIs may be needed to replicate this phenomenon, although it may then become challenging to make AAV preparations of sufficient volume and concentration to achieve such higher MOIs. Moreover, *in vivo* delivery of high doses of AAV has been reported to lead to acute toxicity.^55^

In summary, our findings suggest that, like AAV6, clade F AAVs are unable to mediate high-frequency HDR in the absence of a targeted DNA break. We observed this at 2 different loci in 3 distinct cell types, including primary human HSPCs. While our data agrees that delivery of the donor DNA template is a critical determinant of the rate of genome editing, inclusion of a matched targeted nuclease was required for any detectable site-specific gene insertion. The gene insertion rates we report without ZFNs align with the consensus in the field that AAV vectors typically achieve less than 1% gene insertion in the absence of a catalytic DNA break.^24–26^ At present we are unable to explain our inability to reproduce the findings of Smith *et al.*^40^ Perhaps there are unidentified features in their AAV constructs, or some aspect of their vector production, purification, or titration methods that contributed to this phenomenon. Nevertheless, the reported ability of clade F AAVs to perform highly efficient nuclease-independent genome editing by homologous recombination is clearly not a universal phenomenon.

## Materials & Methods

### AAV plasmids

The AAV6 capsid plasmid pRC6 was purchased from Cell BioLabs (San Diego, CA). AAV9 capsid plasmid p5E18-VD2/9 was obtained from the University of Pennsylvania Vector Core. AAV9-G505R was generated by PCR mutagenesis of p5E18-VD2/9 using the In-Fusion Cloning HD Plus System (Takara Bio, Mountain View, CA). The pITR-CCR5-PGK-GFP vector genome plasmid contains AAV2 ITRs, CCR5 homology arms of 473 bp (left) and 1431 bp (right), a hPGK promoter, eGFP, and a BGH polyA signal has been previously described.^5^ Plasmid pITR-AAVS1 contains homology arms of 801 bp (left), and 568 bp (right),^5^ and a splice acceptor, 2A peptide from thosea asigna virus, eGFP, and BGH polyA (SA-2A-GFP) was synthesized (Genewiz, La Jolla, CA) and inserted into the pAAV-AAVS1 backbone by In-Fusion Cloning to create vector genome pITR-AAVS1-SA-2A-GFP.

### AAV vectors

AAV6 and AAV9 vectors expressing CMV-GFP were purchased from Vigene Biosciences (Rockville, MD). AAV6-CCR5-PGK-GFP was produced at Sangamo Therapeutics as previously described.^5^ All other AAV vectors were produced in-house using the AAV helper free packaging system (Cell Biolabs). Briefly, one day before transfection, 9 × 10^6^ 293AAV cells (Cell Biolabs) were seeded in 15 cm diameter dishes to achieve 70-80% confluence on the day of transfection. Cells were co-transfected with the following three plasmids: pHelper; capsid plasmid pAAV-RC6 for AAV6, p5E18-VD2/9 for AAV9, or p5E18-VD2/9-G505R for AAV9-G505R; and pITR-CCR5-PGK-GFP or pITR-AAVS1-SA-2A-GFP. A total of 81 µg of plasmids were transfected by calcium phosphate transfection method at a 1:1:1 ratio. Sixteen hours after transfection, cells were washed once with PBS and kept in fresh DMEM supplemented with 10% FBS.

For AAV9 and AAV9-G505R production, cell pellets were harvested at 72 hours post-transfection and freeze/thawed at least once with a −80°C/37°C cycle. The cell pellets were then lysed (150mM NaCl, 20mM Tris pH8, 0.5% sodium deoxycholate, and 100 U/mL benzonase) for 2 hours. Cell debris were removed by centrifugation at 3000 g for 15 mins. The crude lysate was subjected to iodixanol gradient ultracentrifugation at 59000 rpm for 70 minutes. After ultracentrifugation, the 40% iodixanol fraction containing AAV vectors was isolated, and further concentrated and buffer exchanged to D-sorbitol containing PBS (PBS with 5% D-sorbitol and 350mM NaCl) by centrifugation through an Amicon Ultra-50 centrifugal filter (Millipore, Burlington, MA) according to the manufacturer’s instructions. The concentrated vectors were stored at −80°C until use.

For AAV6 production, culture medium was harvested at 72 hours post-transfection and filtered through a 0.22 µM filter. The filtered medium was then concentrated 20-fold by tangential flow filtration (TFF) system (Spectrum Laboratories, Rancho Dominguez, CA), using a polyethersulfone membrane hollow fiber unit with 100 kDa molecular weight cut off and 155 cm^2^ filtration surface. The KR2i peristaltic pump was used to pump the medium through the filter. The concentrated medium was then subjected to iodixanol gradient ultracentrifugation as described above.

### AAV vector titration

To remove residual plasmid DNA, AAV vectors were treated with DNaseI (New England Biolabs, Ipswich, MA) at 37°C for 30 mins, followed by heat inactivation at 75°C for 10 mins. DNA was extracted by treatment with proteinase K (Sigma-Aldrich, St. Louis, MO) for 1 hour at 37°C, followed by heat inactivation at 95°C for 20 mins. The extracted DNA was stored at −20°C until titration.

AAV vector genome (vg) titers were determined by TaqMan qPCR (Thermo Fisher, Waltham, MA) using ITR specific primers (AAV ITR-Forward 5’-GAACCCCTAGTGATGGAGTT-3’, AAV ITR-Reverse 5’-CGGCCTCAGTGAGCGA-3’) and probe (AAV ITR-Probe 5’-FAM-CACTCCCTCTCTGCGCGCTCG-Tamra-3’). To prepare the standard curve, serial dilutions of DNA extracted contemporaneously from a recombinant AAV2 Reference Standard Material (American Type Culture Collection, Manassas, VA; VR-1616) was used.^53^

### ZFN reagents and mRNA production

ZFNs targeting the CCR5 and AAVS1 loci have been described previously.^5^ The CCR5-specific ZFNs were used in a bicistronic cassette with a 2A peptide from thosea asigna virus, while AAVS1-specific ZFNs were used as 2 separate monomers. Plasmid DNAs were linearized by restriction enzyme digest (SpeI) and purified using the Zymo DNA Clean and Concentrator protocol (Zymo Research, Irvine, CA). *In vitro* transcription of mRNA was performed using the T7 mScript Standard mRNA Production System (Cellscript, Madison, WI) per manufacturer’s instructions. RNA was purified using RNA Clean & Concentrator-25 (Zymo Research) per manufacturer’s protocol and stored at −80° C until use.

### Isolation and culture of human CD34^+^ HSPCs

Fetal liver CD34^+^ HSPCs were isolated from tissue obtained from Advanced Bioscience Resources (Alameda, CA) as anonymous waste samples, with approval of the University of Southern California’s Institutional Review Board. CD34^+^ cells were isolated as previously described,^5^ using physical disruption, incubation in collagenase to give single cell suspensions, and magnetic bead selection using an EasySep™ Human CD34 Positive Selection Kit (STEMCELL Technologies Inc., Vancouver, BC, Canada). The resulting CD34^+^ HSPCs were cultured in StemSpan SFEM II (STEMCELL Technologies Inc.) supplemented with 1% penicillin/streptomycin/amphotericin B (PSA) (Sigma Aldrich) and SFT cytokines: 50 ng/mL each of SCF, Flt3 ligand and TPO (R&D Systems, Minneapolis, MN).

### Cell culture, AAV transduction, and electroporation of cells

HEK-293T cells and HeLa cells were cultured in DMEM supplemented with 10% FBS and 1% penicillin/streptomycin. Cells were seeded overnight to adhere to plates and washed once with PBS prior to AAV transduction in DMEM without addition of FBS. AAV was added to cells at indicated MOIs, and after 4 h FBS was restored to the culture.

K562 cells were cultured in RPMI-1640 supplemented with 10% FBS and 1% penicillin/streptomycin. For genome editing, cells were washed twice with PBS and resuspended at 2 × 10^7^ cells/mL in RPMI without FBS. Cells were transduced with AAV at 10^4^ vg/cell for 4 h. AAV only samples were diluted to 4 × 10^5^ cells/mL in RPMI with 10% FBS added. For cells also receiving ZFN electroporation, 10 µL of transduced cells were mixed with 90 µL of SF buffer and ZFN mRNA and electroporated using a 4D-X nucleofector using pulse code FF-120 per manufacturer’s recommendation (Lonza, Basel, Switzerland). Cells received 1.6 µg of CCR5-specific ZFN mRNA or 0.7 µg of each AAVS1-specific ZFN mRNA.

After overnight pre-stimulation, HSPCs were washed twice with PBS and resuspended at 1 × 10^7^ cells/mL in BTXpress Electroporation Buffer, mixed with ZFN mRNA, and electroporated using a BTX ECM 830 (Harvard Apparatus, Holliston, MA) at 250 V for 5 ms. Cells were resuspended in SFEM-II with SFT and PSA and transduced with indicated AAV vectors. After 4 h, media was supplemented with 10% FBS and cells were cultured for downstream analyses by flow cytometry.

GFP expression was measured by flow cytometry as indicated on either a FACSCanto II (BD Biosciences, San Jose, CA) or Guava easyCyte (MilliporeSigma, Burlington, MA). Data were analyzed using FlowJo software (Flowjo LLC, Ashland, Oregon).

### Vector copy number analysis

Cells were pelleted by centrifugation and DNA was isolated using the DNeasy Blood & Tissue Kit (Qiagen, Hilden, Germany). Roughly 5 ng of genomic DNA was mixed with primers and probes for a human RPP30 copy number assay labeled with HEX (Bio-Rad, Hercules, CA) as an internal control, and GFP-specific primers and probe were as follows: Forward 5’-AGCAAAGACCCCAACGAGAA-3’, Reverse 5’-GGCGGCGGTCACGAA-3’, Probe 5’-FAM-CGCGATCACATGGTCCTGCTGG-3’. Droplets were prepared using ddPCR Supermix for Probes (No dUTP) and a QX200 Droplet Generator (Bio-Rad). The PCR reaction was run on a C1000 Touch Thermal Cycler with the following conditions: 95 °C 10 min, 40 cycles (94° C 30 s, 60° C 1 min), 98° C 10 min, and 4° C forever. After the PCR reaction, the samples were read on a QX200 Droplet Reader and the data were analyzed with QuantaSoft analysis software (Bio-Rad). Linear regression analysis was performed using GraphPad Prism software (San Diego, CA).

### In-out PCR

Cells were pelleted by centrifugation and DNA was isolated using the DNeasy Blood & Tissue Kit (Qiagen). PCR reactions were prepared with 200 ng genomic DNA and AmpliTaq Gold 360 Master Mix (Applied Biosystems, Foster City, CA). Primers were as follows: CCR5-in 5’-GAGGATTGGGAAGACAATAGCAG-3’, CCR5-out 5’-CCAGCAATAGATGATCCAACTCAAATTCC-3’, AAVS1-in 5’-CTAGGGCCGGGATTCTCCT-3’, AAVS1-out 5’-CGGAACTCTGCCCTCTAACG-3’. Thermal cycling for CCR5 was as follows: 95° C for 10 mins, 35 cycles (95° C 30 s, 55° C 30 s, 72°C 105 s), 72° C 7 mins, and 4° C forever. AAVS1 cycling was identical except the 72° C extension was performed for 60 s. Equal volumes of PCR reactions were run on a 1% agarose gel and visualized with GelRed Nucleic Acid Stain (Biotium, Fremont, CA).

## Acknowledgements

This work was supported by National Institutes of Health grant HL129902 to P.M.C. and a Taiwan USC scholarship to H-Y.C.

## Author contributions

G.L.R., H.Y.C., and H.M. performed experiments. G.L.R. and P.M.C. designed experiments, analyzed and interpreted data, and wrote the manuscript. P.M.C. supervised the study.

